# Lineage structure of *Streptococcus pneumoniae* is driven by immune selection on the groEL heat-shock protein

**DOI:** 10.1101/082990

**Authors:** José Lourenço, Eleanor R. Watkins, Uri Obolski, Samuel J. Peacock, Callum Morris, Martin C. J. Maiden, Sunetra Gupta

## Abstract

Populations of *Streptococcus pneumoniae* (SP) are typically structured into groups of closely related organisms or lineages, but it is not clear whether they are maintained by selection or neutral processes. Here, we attempt to address this question by applying a machine learning technique to SP whole genomes. Our results indicate that lineages evolved through immune selection on the groEL chaperone protein. The groEL protein is part of the *groESL* operon and enables a large range of proteins to fold correctly within the physical environment of the nasopharynx, thereby explaining why lineage structure is so stable within SP despite high levels of genetic transfer. SP is also antigenically diverse, exhibiting a variety of distinct capsular serotypes. Associations exist between lineage and capsular serotype but these can be easily perturbed, such as by vaccination. Overall, our analyses indicate that the evolution of SP can be conceptualized as the rearrangement of modular functional units occurring on several different timescales under different pressures: some patterns have locked in early (such as the epistatic interactions between *groESL* and a constellation of other genes) and preserve the differentiation of lineages, while others (such as the associations between capsular serotype and lineage) remain in continuous flux.

## Introduction

Streptococcus pneumoniae (the pneumococcus) is a gram-positive bacterial pathogen which, although commonly carried asymptomatically in the nasopharynx, can cause pneumonia, meningitis, septicemia and bacteremia in the young, elderly and immuno-compromised, being responsible for about 11% of worldwide deaths in children under 5 years of age^1,2^. Populations of *S. pneumoniae* are antigenically diverse and can be stratified into more than 90 serotypes according to the antigenic properties of the expressed polysaccharide capsule, of which only 10-15 are responsible for most cases of invasive disease worldwide^3^. Reductions in disease rates have been achieved by the deployment of the PCV7 vaccine targeting 7 of the most common serotypes in invasive disease, and more recently through the use of PCV13 which extends coverage to an additional 6 serotypes. However this has been accompanied by an increase in the frequency of non-vaccine serotypes in many parts of the world, likely due to the removal of competition from vaccine serotypes^4^.

Like many other bacterial pathogen populations, *S. pneumoniae* may be organised into a number of so-called clonal complexes on the basis of allelic diversity at selected housekeeping loci (determining Multilocus Sequence Type,^5, 6^). Pneumococcal populations are also structured at a whole genome level into co-circulating lineages or Sequence Clusters (SC) bearing unique signatures of alleles^7–9^. The relationships between clonal complex, lineage and serotype are often found to be non-overlapping^8, 10^, although subject to perturbations such as through vaccination^11^.

The maintenance of discrete major lineages, and their associations with distinct serotypes and clonal complexes, is hard to ascribe to purely neutral processes, given the high rate of genetic exchange in these pathogen populations^12, 13^. We have previously proposed that extensive co-adaptation between loci may give rise to these patterns, as even small fitness differences among different combinations of alleles can lead to the loss of less fit genotypes under intense competition for resources^14^. Bacterial populations could also segregate into a set of successful *metabolic types* which are able to co-circulate by virtue of exploiting separate metabolic niches and thereby avoiding direct resource competition^15^. As an example, specific differences in the ability to absorb particular carbohydrate resources have been observed in functional genomics studies of *S. pneumoniae* ^16^, and these may reflect specialization upon different resources within the same environment as a means of avoiding competition. Establishing the contribution of co-adaptation and competition in the maintenance of discrete lineages is important since the outcome of certain interventions, such as vaccination, depends crucially on these underlying determinants of population structure^17^.

Here, we attempt to elucidate potential drivers of lineage structure by applying a machine learning technique known as the Random Forest Algorithm (RFA) to a dataset containing 616 whole genomes of *S. pneumoniae* collected in Massachusetts (USA) between 2001 and 2007^8^. RFA-based methods have been robustly applied in genome-wide association studies of cancer and chronic disease risk^18^, species classification^19^, or in the search of viral determinants for host tropism, for instance by identifying the key amino-acid sites that determine host specificity of zoonotic viruses^20^, and by selecting the clear genetic distinctions in avian and human proteins of Influenza viruses^21, 22^. In the context of bacterial pathogens, these methods have been sucessfully used to analyse the genetic background of *Escherichia coli* cattle strains more likely to be virulent to humans^23^, to identify *Staphylococcus aureus* genetic variants associated with antibiotic resistance^24^, and to discover that repertoires of virulence proteins within different Legionella species are largely unique (non-overlapping)^25^. An RFA is an ensemble method that combines the information of multiple, regression or classification trees built around predictor variables towards a response variable. The output of an RFA is composed both of the classification success rates of the response variable and a ranking of the predictor variables quantifying their relative role in the classification process.

We used as response variables (i) the capsular serotype of each isolate (which had been determined by serological means), and (ii) the monophyletic Sequence Cluster (SC) to which samples had been assigned^8^. We set the predictor variables to be the 2135 genes for which we had obtained allelic profiles (effectively using a whole-genome multi-locus sequence typing approach^26^) for each of the 616 isolates^17^. Using this method, we confirmed that capsular genes predict serotype, but found a clear disjunction between these genes and those which predict SC (lineage). Furthermore, our analyses revealed that, contrary to the expectations of neutrality, genes which predict lineage are non-randomly distributed across the genome, clustering within and around the *groESL* operon, leading us to propose that a combination of immune selection and coadaptation operating upon these loci may be the primary determinants of lineage structure.

## Results

### Classification success for serotype and sequence cluster

Classification of SC by the RFA was accurate (Fig. S1B) with all SC types being predicted with success close to 100%. This is a reflection of the strong correspondence between classification trees and taxonomy when based on genetic information, as explored in other studies^19^, and demonstrated by Austerlitz and colleagues when comparing the success of RFA, neighbour-joining and maximum-likelihood methodologies on simulated and empirical genetic data^27^.

By contrast, the success rate in identifying the capsular serotypes of the 616 whole genomes, although also very high (above 75% for the majority of serotypes), was not perfect (Fig. S1A). This is to be expected given the imperfect association between lineage and serotype, and also because certain serotypes were represented by very small numbers of isolates (as an extreme example, only a single isolate of serotype 21 was present and therefore classification success was nil).

### The capsular locus is a strong predictor of serotype but performs indifferently in predicting sequence cluster

As might be expected, genes within the capsular locus (defined as being within but not including the genes *dexB* and *aliA*) were higly predictive of serotype, with their RFA scores appearing as outliers in the top 2.5% of the distribution defined by all 2135 genes in the dataset (Fig. 1A, see Methods). However, these did not score above average in predicting SC, as their RFA scores shifted closely to the distribution’s average (Fig. 1B). We noted, however, that many of these genes contained what appeared to be a high proportion of deletions but, in fact, had simply eluded allelic notation (alleles ’x’, see Methods) on account of their high diversity at the level of the population (see, for example,^28^). For certain genes, such as those encoding the polysaccharide polymerase Wzy and the flippase Wzx, the allelic notation process failed at least 50% of the time for over 90% of the isolates, essentially working only for serotype 23F (the reference genome) and the closely related 23A and 23B serotypes. In general, the degree of success in allelic notation of each gene was closely linked to the potential for alignment with its counterpart in the 23F reference genome (Fig. S4). Nonetheless, the same shift towards lower RFA scores of capsule-associated genes in predicting SC rather than serotype was observed upon performing a series of sensitivity classification exercises after excluding all genes which contained *>* 50% (Fig. S2) or *>* 10% (Fig. S3) of gene mismatches/deletions. When imposing an exclusion criterion of *>* 10% we retained only the genes *wze*, *wzg* and *wzh* (in addition to two pseudogenes) within the capsular locus, and these could also clearly be seen to shift from above the upper 97.5% limit into the neutral expectation of RFA scores when predicting SC (Fig. S3).

**Figure 1.**
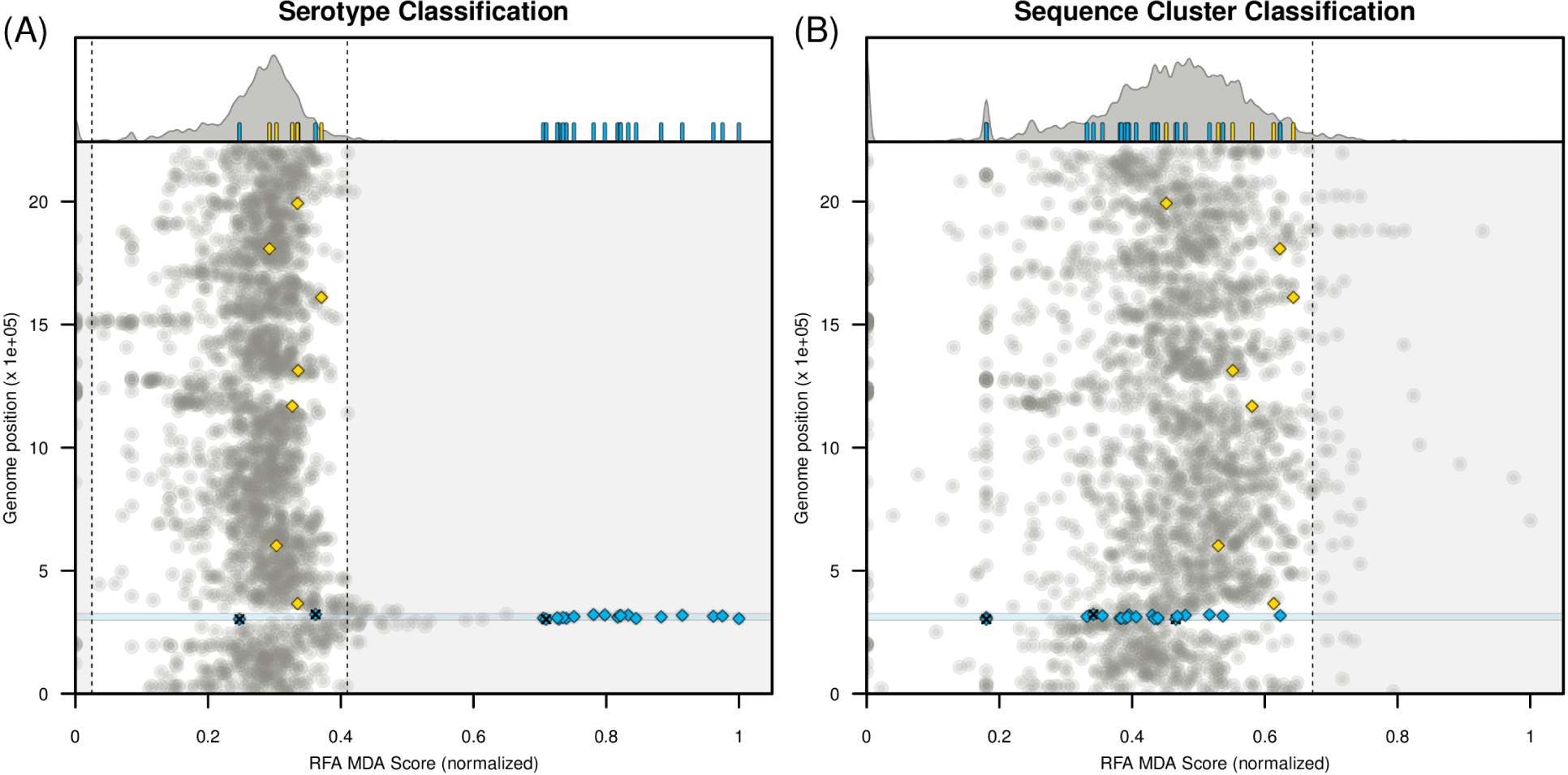
Random forest classification. *(A)* Random forest analysis (RFA) for serotype classification. *(A, top)* Density function of RFA scores obtained for each gene in the dataset. The 95% boundaries are marked by the dashed lines. Small bars highlight the RFA scores of genes within particular groups (yellow for MLST genes, blue for capsular locus genes). *(A, bottom)* Genomic position for each gene in the dataset against their RFA score (normalised to [0,1]). The circular genome is presented in a linear form on the y-axis, with the first gene being *dnaA* and the last gene *parB*. MLST genes are marked in yellow diamonds (*spi, xpt, glkA, aroE, ddlA, tkt*) and genes within the capsular locus with blue diamonds (pseudogenes tagged with ’x’). *(B)* RFA analysis for sequence cluster classification; figure details the same as in *A*. Blue shaded areas in both A and B subplots mark the capsular locus (genes within *aliA* and *dexB*).

We next performed the same analysis excluding all genes which showed mismatches or deletions above a threshold of 1%, in an attempt to eliminate possible biases in RFA output due residual information arising from the distribution of mismatches/deletions. This left us with 1581 genes which were shared by virtually all the samples in our dataset and for which function could be correctly ascertained by querying the reference genome. It is likely that these genes correspond to the approximately 1500 core cluster of orthologous genes (COGs) identified by Croucher *et al* in their recent analysis of the same dataset^7^. This approach eliminated all of the genes considered above as belonging within the capsular locus, although flanking genes were retained and a number of these achieved the top 2.5% of RFA scores in predicting serotype (Fig. 2A, Table 1): 38% of the top genes occurred within 10 genes downstream and upstream of the capsular locus, and 90% were situated within 129 genes (which amounts for 6% of the genome). The remaining 10% of top-scoring genes, *lytC*, *trpF*, *patB* and SPNF2300400 were located at significantly longer distances from the capsular locus, at 963 (45% of the genome), 710, 469 and 270 (13% of the genome) genes away, respectively. None of the genes achieving the top 2.5% of RFA scores in predicting serotype (shown in red in Fig. 2) remained in the top 2.5% category when asked to predict SC. Similarly, all genes which achieved top scores in predicting SC (Table 2) were only of average importance in elucidating serotype (shown in green in Fig. 2).

**Table 1.**
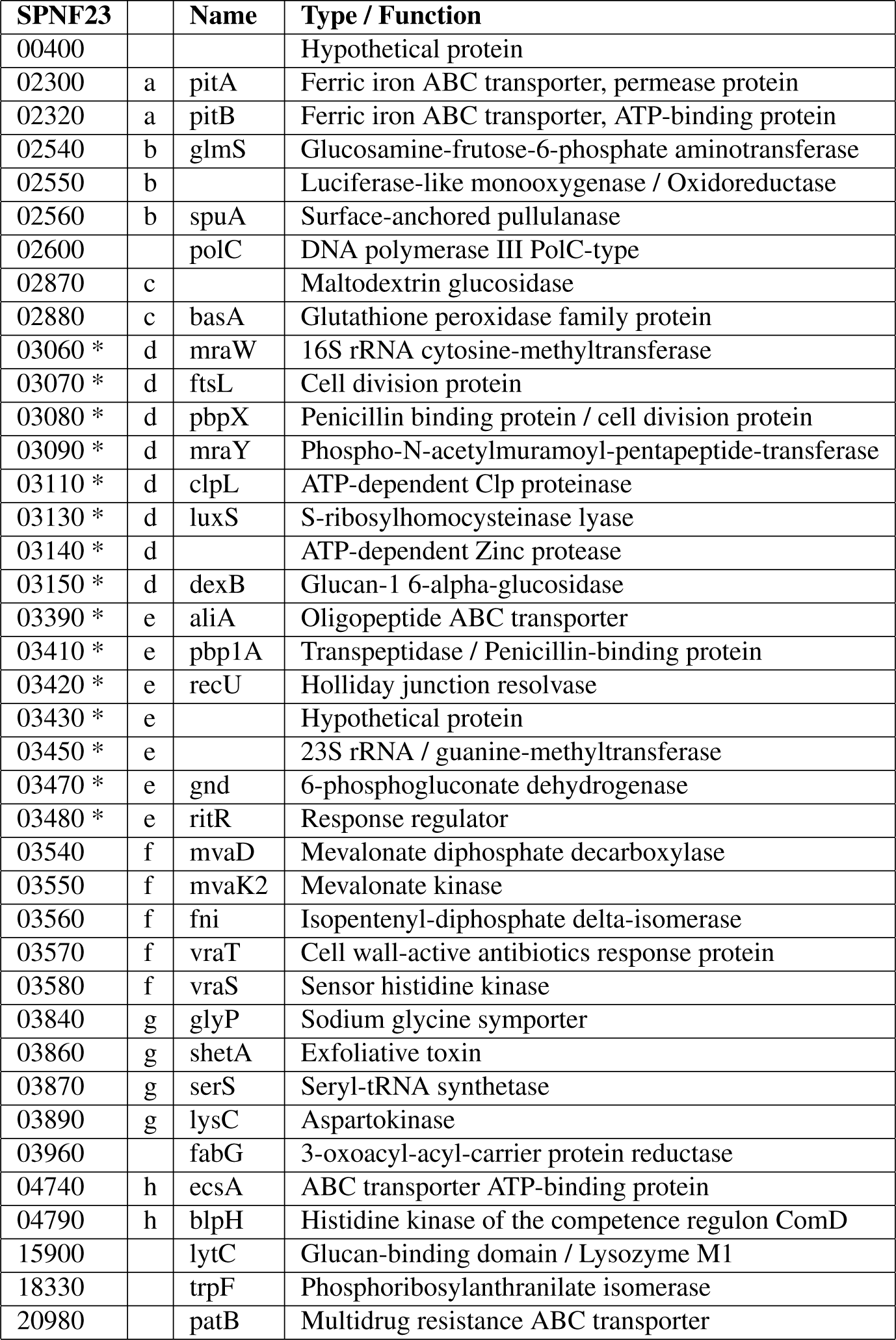
Top genes for Serotype prediction. Genes marked with * flank up to 10 genes, upstream or downstream from the capsular locus. Letters *a* to *h* denote groups of contiguous genes (minimum proximity of 2 genes).

| SPNF23 |  | Name | Type / Function |
| --- | --- | --- | --- |
| 00400 |  |  | Hypothetical protein |
| 02300 | a | pitA | Ferric iron ABC transporter, permease protein |
| 02320 | a | pitB | Ferric iron ABC transporter, ATP-binding protein |
| 02540 | b | glmS | Glucosamine-fructose-6-phosphate aminotransferase |
| 02550 | b |  | Luciferase-like monooxygenase / Oxidoreductase |
| 02560 | b | spuA | Surface-anchored pullulanase |
| 02600 |  | polC | DNA polymerase III PolC-type |
| 02870 | c |  | Maltodextrin glucosidase |
| 02880 | c | basA | Glutathione peroxidase family protein |
| 03060 * | d | mraW | 16S rRNA cytosine-methyltransferase |
| 03070 * | d | ftsL | Cell division protein |
| 03080 * | d | pbpX | Penicillin binding protein / cell division protein |
| 03090 * | d | mraY | Phospho-N-acetylmuramoyl-pentapeptide-transferase |
| 03110 * | d | clpL | ATP-dependent Clp proteinase |
| 03130 * | d | luxS | S-ribosylhomocysteinase lyase |
| 03140 * | d |  | ATP-dependent Zinc protease |
| 03150 * | d | dexB | Glucan-1 6-alpha-glucosidase |
| 03390 * | e | aliA | Oligopeptide ABC transporter |
| 03410 * | e | pbp1A | Transpeptidase / Penicillin-binding protein |
| 03420 * | e | recU | Holliday junction resolvase |
| 03430 * | e |  | Hypothetical protein |
| 03450 * | e |  | 23S rRNA / guanine-methyltransferase |
| 03470 * | e | gnd | 6-phosphogluconate dehydrogenase |
| 03480 * | e | ritR | Response regulator |
| 03540 | f | mvaD | Mevalonate diphosphate decarboxylase |
| 03550 | f | mvaK2 | Mevalonate kinase |
| 03560 | f | fni | Isopentenyl-diphosphate delta-isomerase |
| 03570 | f | vraT | Cell wall-active antibiotics response protein |
| 03580 | f | vraS | Sensor histidine kinase |
| 03840 | g | glyP | Sodium glycine symporter |
| 03860 | g | shetA | Exfoliative toxin |
| 03870 | g | serS | Seryl-tRNA synthetase |
| 03890 | g | lysC | Aspartokinase |
| 03960 |  | fabG | 3-oxoacyl-acyl-carrier protein reductase |
| 04740 | h | ecsA | ABC transporter ATP-binding protein |
| 04790 | h | blpH | Histidine kinase of the competence regulon ComD |
| 15900 |  | lytC | Glucan-binding domain / Lysozyme M1 |
| 18330 |  | trpF | Phosphoribosylanthranilate isomerase |
| 20980 |  | patB | Multidrug resistance ABC transporter |

**Table 2.**
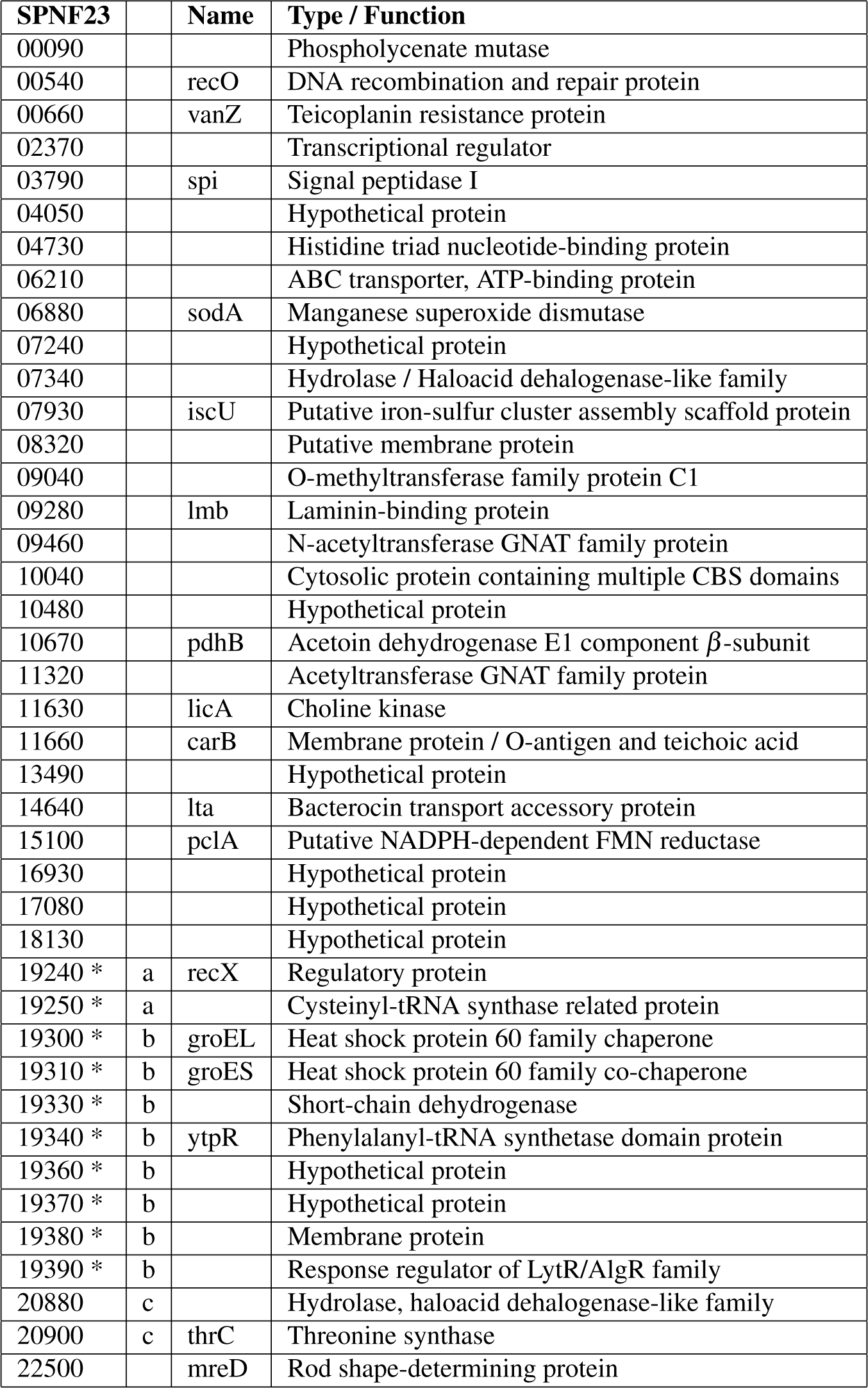
Top genes for Sequence Cluster prediction. Genes marked with * flank up to 10 genes, upstream or downstream from the *groESL* operon. Letters *a* to *c* denote groups of contiguous genes (minimum proximity of 2 genes).

### Sequence cluster is predicted by variation in the groESL operon

About 75% of top-scoring genes for SC were uniformly distributed along the genome (Fig. 2B), while the remaining 25% of the genes were contiguous and contained within the *groESL* operon, encoding the GroEL chaperone and GroES co-chaperone proteins (Table 2). Notably, other studies have reported the power of the *groESL* operon and its proteins to ascertain phylogeny and classification within the *Streptococcus* genus^29^ and between species of the *Viridans* and *Mutans* Streptococci groups^30, 31^.

A number of other top scoring genes in predicting SC have also previously been demonstrated to be powerful discriminators of genealogy in a range of bacterial species. For instance, *sodA*, encoding for the manganese superoxide dismutase, critical against oxidative stress and linked to both survival and virulence, has been highlighted in numerous studies for its relevance in identification of rare clones of pneumococci^32, 33^ and Streptococci at the species level^34, 35^. Another example is the *lmb* gene, encoding for an extracellular protein with a key role in physiology and pathogenicity^36, 37^. Homologs of this protein have been documented to be present and discriminatory of at least 25 groups of the *Streptococcus* genus with possible similar functions^38, 39^.

The housekeeping genes included in multilocus sequence typing (MLST) classification performed no better than average in predicting SC across the sensitivity experiments (Fig. 1, S1-3). The exception was the Signal Peptidase I gene (*spi*), which featured in the top-scoring genes predicting SC under the strict 1% cutoff (Table 2). This is unsurprising, however, as MLST genes are unlikely to dictate lineage differentiation through selective processes, which endorses their choice as good discriminators of recent neutral diversification, in particular within recent epidemiological events^5, 6^.

**Figure 2.**
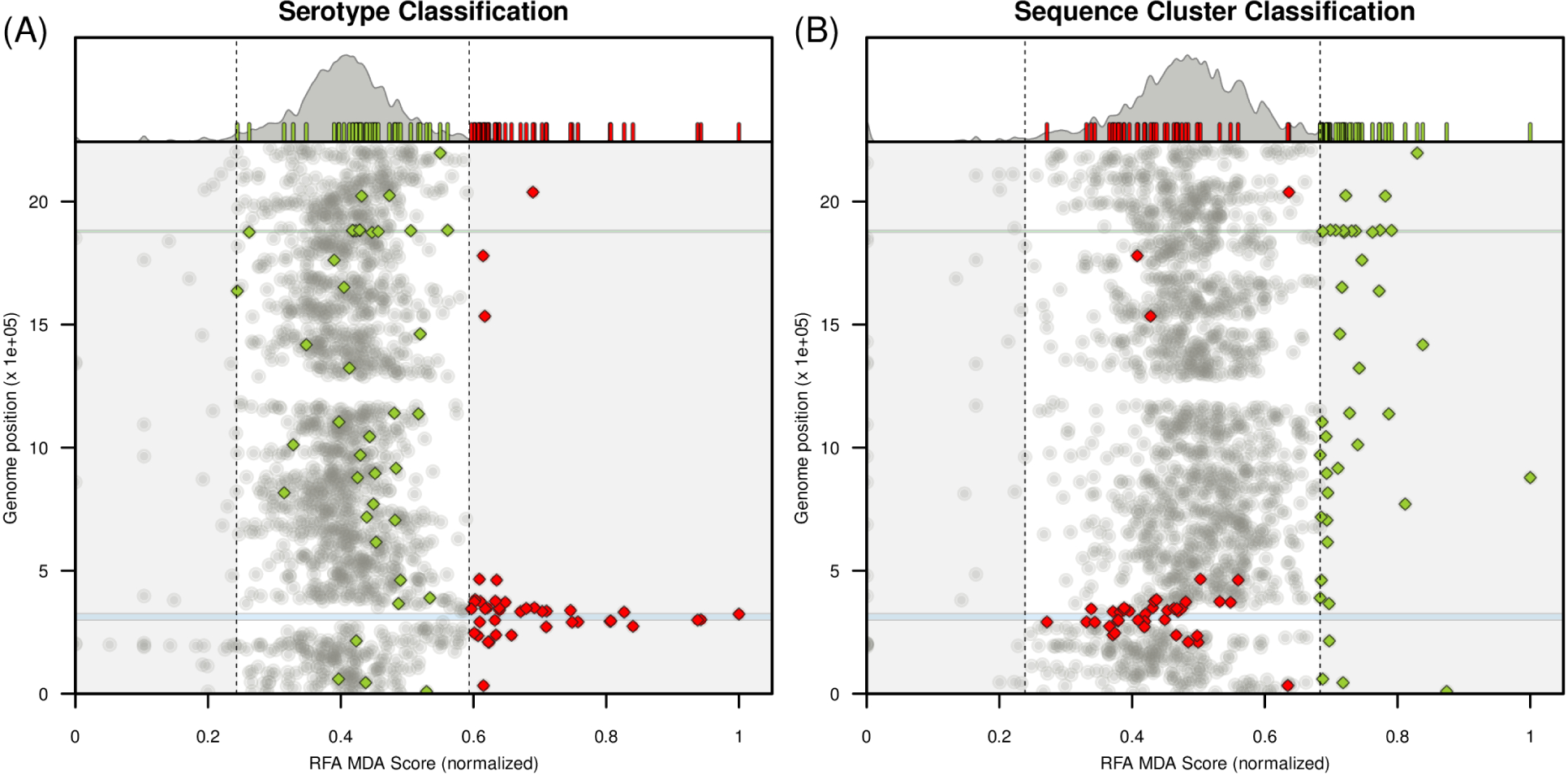
Random forest classification excluding data with gene mismatches. *(A)* Random forest analysis (RFA) for serotype classification when excluding genes for which the allelic notation process had *<* 99% positive matches with the reference genome. *(A, top)* Density function of RFA scores obtained for each gene in the dataset. The 95% boundaries are marked by the dashed lines. Small bars highlight the RFA scores of genes within particular groups (red for serotype, green for SC genes). *(A, bottom)* Genomic position for each gene in the dataset against their RFA score (normalised to [0,1]). The circular genome is presented in a linear form on the y-axis, with the first gene being *dnaA* and the last gene *parB*. Red and green diamonds mark the top 2.5% ranking genes for serotype and Sequence Cluster classification, respectively. *(B)* RFA for Sequence Cluster classification; figure details the same as in *A*. Blue shaded areas in both A and B mark the capsular locus (genes within *aliA* and *dexB*). Green shaded areas in both A and B mark the genes contiguous and including the *groESL* operon (Table 1).

### Top-scoring genes for serotype are associated with resource competition and antibiotic resistance

When analyzing the 39 top-scoring, non-capsular genes which were highly predictive of serotype, we found 24 (62%) with compelling support for functional background that could mediate pneumococcal competitive interactions or niche specialization, at least in related streptococcal species (reviewed in detail in supplementary text). For instance, ATP-binding cassette (ABC) transporter genes, critical for intake, antibiotic resistance and metabolism, were found 5 times more frequently in the genes predictive of serotype compared to those determining SC (Tables 1 and 2). Notably, our approach selected the genes encoding for the pit ABC transporter, a key player in iron uptake known to exhibit strain-specific variation^40^, but did not select two other operons encoding iron transporters (*piu*, *pia*), which are conserved between *S. pneumoniae* strains^40^ and therefore unlikely to be predictors of serotype. Transport of essential substrates is also achieved by alternative systems which were also captured by our approach, such as the passive channel sodium symporter GlyP^41^ or the use of menaquinones and ubiquinones for electron transport (mevalonate pathway)^42–44^. We also found some of the top-scoring entries to be involved in functions associated with respiration (*ecsA, mvaD, mvaK2*) and amino acid, fatty acid and cell wall or capsular biosynthesis which amounted for approximately 25% of the top-scoring genes (*trpF, fabG, lysC, mvaD, mvaK2, ritR, pbp1A, pbpX, mraW* and *mraY*).

High RFA scores for serotype were also found among a number of genes flanking the capsular locus which are involved in antibiotic resistance, such as penicillin-binding protein genes *pbpX* and *pbp1A*, the 16S rRNA cytosine-methyltransferase gene *mraW* and the phospho-N-acetylmuramoyl-pentapeptide-transferase gene *mraY*. Genes involved in resistance to other antibiotics such as tomethicillin, vancomycin, daptomycin (*vra* operon)^45^ and the broad-spectrum quinolones family (*patB*)^46–48^were also featured in the top-scoring genes. Also of note were entries linked to direct inter- and intra-species competition, either through factors related to immune escape or warfare. These included genes linked to pneumolysin expression and biofilm formation (*luxS*)^49, 50^, and production of bacteriocins (*blpH*)^51, 52^, ammonia (*glmS*)^53^ and lysozymes (*lytC*)^54, 55^.

### Several top-scoring genes for SC classification are also key determinants of phenotype

The top-scoring genes predicting SC were discordant to the ones determining serotype and approximately 30% were found to have unknown functions (Table 2). However, we also found several examples of genes whose functions (reviewed in supplementary text) would be expected to be naturally linked with particular phenotypes such as virulence (*sodA, lmb, pdhB, varZ, licA*)^32, 33, 36, 37, 56–58^ or specific virulence traits such as host-cell adherence (*pclA*)^59^ or laminin binding (*lmb*)^39^. Several genes were also found to encode or directly produce proteins or protein-complexes which are highly immunogenic, such as the *groEL*^60–62^, *lmb* ^39^, *carB* ^63, 64^, and *licA* ^58, 65^ genes.

## Discussion

Our aim, in this paper, was to test the hypothesis that the stratification of pneumococcal populations into distinct sequence clusters or “lineages” occurred through neutral processes, with serotype diversity being superimposed upon the ensuing “clonal” framework to minimize antigenic interference between lineages. To this end, we applied a Random Forest Algorithm (RFA) to assess the contribution of different genes in determining the serotype or sequence cluster of isolates within a dataset containing 616 whole genomes of *S. pneumoniae* collected in Massachusetts (USA)^8^, for each of which we had obtained allelic profiles of 2135 genes of both known and unknown function^17^. By selecting the 2.5% of top RFA scores, we effectively focused on the subset of possibly selected units (genes) which present combinations of alleles that appear statistically more informative than expected at the genome level (see Methods for details). We show that by comparing the genomic localization and function of these top-scoring units (genes), general expectations concerning population structure can be revisited^66^, and inferences can be made concerning the evolutionary processes underlying the formation, relationship and maintenance of serotype and sequence cluster (lineage) at the population level.

Reassuringly, genes of the capsular locus (cps) and many of those flanking it achieved high RFA scores in predicting serotype. We also found a preponderance of genes scoring highly for serotype prediction to be associated with key functions that could define unique *metabolic types* that would have diversified in order to avoid direct resource competition, as previously proposed^4, 17^. Most of these genes were at a distance of less than 6% of the genome to the cps locus. However, linkage disequilibrium is extremely high in this dataset (Fig S7), regardless of genetic distance (with only a slight increase between loci that are less than 10 genes apart); thus, it was not possible to determine whether these had become segregated through competition or by physical and/or functional associations with this locus.

Genes that were highly informative in predicting lineage (sequence cluster) were entirely distinct from those determining serotype. Contrary to what would be expected from a population structure maintained mostly by neutral processes, around a quarter of these genes co-localized within and around the *groESL* operon (marked with * in Table 2), which encodes the macromolecular machinery for a well-studied protein folding system centred around the chaperone GroEL and co-chaperone GroES^67^. In *Escherichia coli*, approximately 10% of total cytosolic proteins, including 67 essential proteins, have been demonstrated to have stable binding to GroEL, with 50 of these confirmed to depend on *groESL* folding via GroEL-depletion experiments^68^. GroEL is also known to be highly immunogenic in *S. pneumoniae* ^61, 69^, as well as in other bacterial species^62, 70^, 71. This raises the radically alternative possibility that sequence clustering may have arisen from immune selection operating on groEL in conjunction with extensive coadaptation with genes encoding the proteins which rely on this chaperonin system.

Our results provide a mechanistic basis for the distinction proposed by Croucher and colleagues^7^, in the context of the same dataset, between infrequent macroevolutionary changes providing a stable backdrop for more frequent, and often transient, microevolutionary changes (see Figure 3). The differentiation of the *groESL* operon would be a striking example of macroevolution, not only driving the emergence of *S. pneumoniae* sequence clusters but also serving to genealogically distinguish closely related bacterial species^29–31^. Several other genes scoring highly for SC were also found to encode or directly produce proteins or protein-complexes which are highly immunogenic (eg. *lmb, carB, licA*), and these may contribute to the maintenance of lineage structure by co-selection with groEL in accordance with the strain theory of host-pathogen systems in which immune selection operating on multiple immunogenic loci can cause the emergence of non-overlapping combinations of alleles^74–76^. In contrast, the emergence and maintenance of serotypes within major lineages (SC) would be dictated by differentiation in genes within and surrounding the capsular locus, and less permanent associations could arise between SC and serotype (at a microevolutionary scale) through resource competition^14, 17^ or indeed multi-locus immune selection operating on GroEL, cps, as well as other surface antigens^73^.

**Figure 3.**
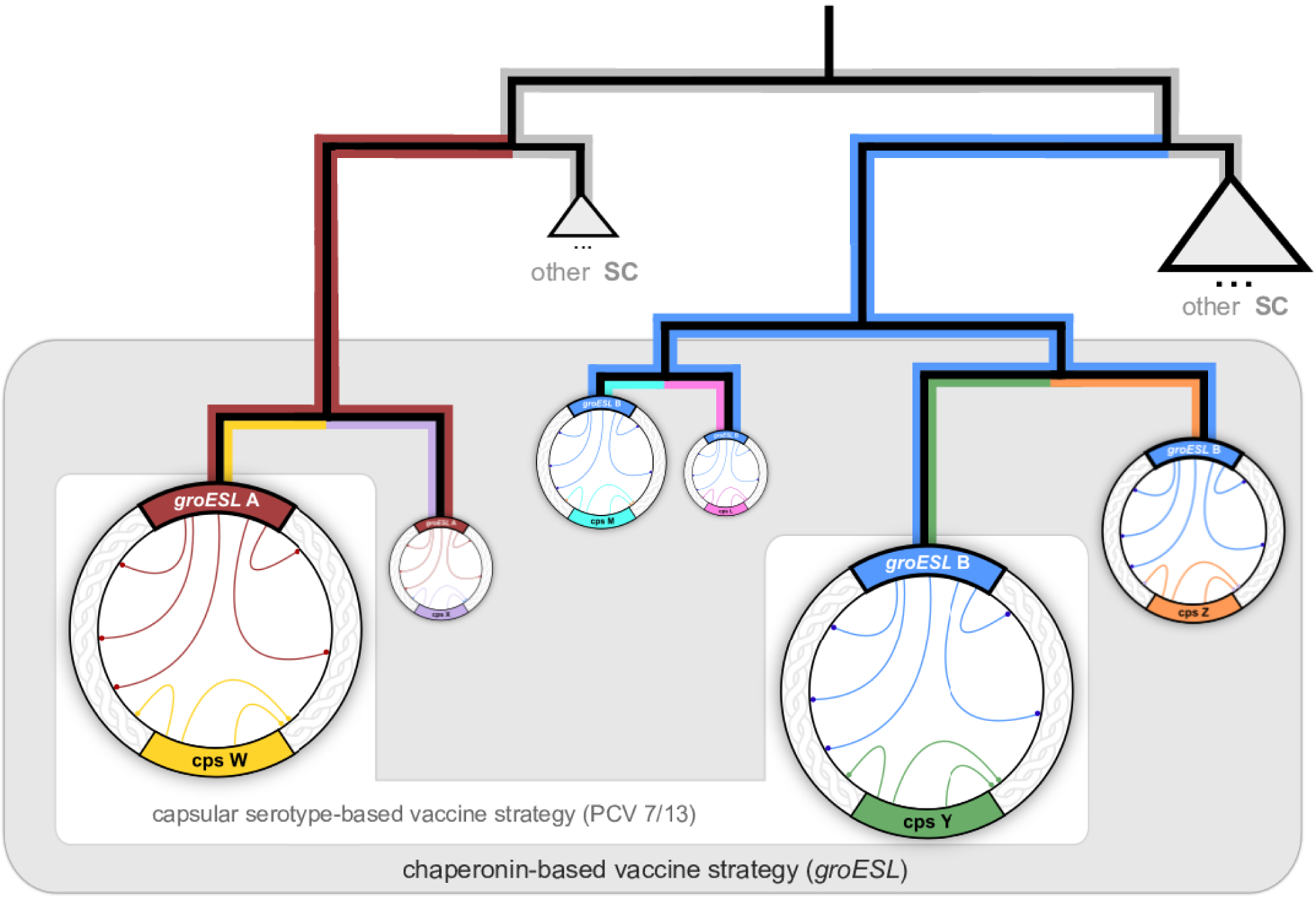
Population structure and vaccination. Conceptual representation of phylogenetic relationships between serotypes and Sequence Clusters (SC), where the former are defined by variation at the cps locus (arbitrarily designated X, W, Y, Z, M, and L, respectively coloured purple, yellow, green, orange, cyan and pink) and the latter are linked to variation in the *groESL* operon (arbitrarily designated A and B and respectively coloured red and blue). Circles symbolize genotypes, with size relative to their prevalence at the population level. Inner genome arcs represent epistatic links: those with the *groESL* operon extend across the genome, while links with the cps locus are more local. Within our framework and according to observed patterns^8^, most SCs will be dominantly associated with a single serotype. Current vaccine strategies (white area) that target a selection of capsular serotypes can lead to the expansion of non-vaccine serotypes (VISR,^72, 73^), potentially within the same sequence cluster (VIMS^17^). Vaccine strategies based on *groESL* variants (grey area) would target entire SCs instead, including all uncommon serotypes within and thereby preventing their expansion.

We note that genes belonging to the Rec family are positioned in close proximity to both the contiguous clusters of top-scoring genes for SC and serotype (Tables 1 and 2). For example, the top-scoring gene *recX* is in close proximity to the *groESL* operon and encodes a regulatory protein that inhibits the RecA recombinase in multiple species of bacteria^77–80^. Restriction-modification systems (RMS) have been proposed as a means of maintaining species identity in a number of bacterial systems^81^ and this idea has been extended to the maintenance of lineages within meningococcal^82^ and pneumococcal^7^ species. Within our framework, RMS would act at even more local scale, principally to conserve the function of critical operons such as *groESL*, rather than prevent their recombination with other genes or operons. It has recently been demonstrated that GroEL in *E. coli* can be functionally replaced, at least partially, by an eukaryotic chaperonin^83^ indicating that the maintenance of particular associations of genes with the *groESL* operon is a consequence of their superior fitness rather than an inability to recombine. We would therefore argue that RMS play a role in protecting the modularity of the genome and that population structure arises through selection favouring particular combinations of variants of these modules.

Overall, our analyses support the hypothesis that lineage structure in maintained by co-adaptation and competition (Buckee et al, 2008) and show, surprisingly, that these selection pressures converge upon the capsular locus and *groESL* operon. Our results endorse the development of vaccines against the associated chaperone protein, groEL, since targetting its protein folding machinery may provide a robust method (Fig. 3) of eliminating particular highly successful lineages rather than promoting the survival of those genotypes within it which carry cps loci encoding non-vaccine capsular serotypes (Watkins et al, 2015). We hope, for these reasons, that this work will stimulate further empirical testing of our hypothesis that immune selection against groEL may be a primary driver of lineage differentiation in the pneumococcus.

## Methods

### Sequence Data and Allelic Annotation

We used a dataset sequenced by Croucher et al, comprising 616 carriage *S. pneumonaie* genomes isolated in 2001, 2004 and 2007 from Massachusetts (USA). The data included 133, 203, 280 samples from 2001, 2004, 2007, respectively; and is stratified into 16 samples of serotype 10A, 50 of 11A, 7 of 14, 24 of 15A, 60 of 15BC, 8 of 16F, 5 of 17F, 6 of 18C, 73 of 19A, 33 of 19F, 1 of 21, 21 of 22F, 33 of 23A, 23 of 23B, 17 of 23F, 11 of 3, 4 of 31, 5 of 33F, 6 of 34, 49 of 35B, 18 of 35F, 2 of 37, 9 of 38, 47 of 6A, 17 of 6B, 33 of 6C, 3 of 7C, 11 of 7F, 4 of 9N, 6 of 9V and 14 of NT (see^8^ for collection details). In summary, we performed a whole genome multi-locus sequence typing (wgMLST) allelic notation^26^ using the BIGSdb software with an automated BLAST process^84^ and the Genome Comparator tool (with ATCC700669 serotype 23F, accession number FM211187, as the reference genome)^17^. This wgMLST approach resulted in the identification of 2135 genes in common between the reference and all the samples in the dataset. Alleles identical to the reference were classified as ’1’, with subsequent sequences, differing at least by one base, labelled in increasing order. Genes were further classified as allele ’X’ when genetic data present had no match to the genome of interest, or were found to be truncated or non-coding. For a visual representation of the allelic annotation and diversity please refer to S1 dataset of^17^. The allelic matrix as obtained by this approach and used in the RFA analysis is herein made available in supplementary Table S1, which also includes the Accession Numbers, gene name, gene product, gene position in reference genome, and year of collection, Sequence Cluster and serotype of each sample.

### Random Forest Approach

We implement a machine learning approach based on a Random Forest Algorithm (RFA) to predict particular features (serotype or Sequence Cluster) of each pneumococci isolate from information on the wgMLST allelic composition of the 2135 genes^85^. In summary, the RFA process takes the following pseudo-steps: (I) the response variable and predictor variables are chosen by the user; (II) a predefined number of independent bootstrap samples are drawn from the dataset with replacement, and a classification tree is fit to each sample containing roughly 2/3 of the data, for which predictor variable selection on each node split in the tree is conducted using only a small random subset of predictor variables; (III) the complete set of trees, one for each bootstrap sample, composes the random forest, from which the status (classification) of the response variable is predicted as an average (majority vote) of the predictions of all trees. Compared to single classification tress, RFs increase prediction accuracy, since the ensemble of slight different classification results adjusts for the instability of the individual trees and avoids data overfitting^86^.

Here we use randomForest: Breiman and Cutler’s Random Forests for Classification and Regression, a software package for the R-statistical environment^87^. Predictor variables are set to be each gene in our genome samples and the response variable is set to the serotype or Sequence Cluster classification of each genome (as per^8^). We use the Mean Decrease Accuracy (MDA), or Breiman-Cutler importance, as a measure of predictor variable importance, for which classification accuracy after data permutation of a predictor variable is subtracted from the accuracy without permutation, and averaged over all trees in the RF to give an importance value^86^. The strategy herein employed is not of quantitative nature, as the absolute scale of scores produced by the RFA is dependent on the dataset being analyzed^85^. Instead, we focus on the 2.5% of top RFA scores as presented by the resulting MDA distribution for all genes, thus selecting the subset of genes which present combinations of alleles that appear statistically more informative than expected at the genome level (i.e. we assume that 95% of the scores should fall between the 2.5th and 97.5th percentiles). With this assumption and the approach detailed below, we effectively select the genes which present a p-value *<* 0.05 given an intrinsic distribution of scores generated by data permutation (a null distribution of scores).

For the results presented in the main text, we assume the predictor variables to be numerical (as opposed to categorical). This assumption is known to introduce RF biases, as classification is effectively made by regression and artificial correlations between allele numbering and the features being selected (serotype and Sequence Cluster) may be present. The assumption is herein necessary since the RFA R-based implementation (version 3.6.12) has an upper limit of 53 categories per predictor variable and we find some genes to present up to 6 times this limit in allele diversity. The categorical constraint is a common feature of RFA implementations, as predictor variables with N categories imply 2*^N^* possible (binary) combinations for an internal node split, making the RFA method computationally impractical. Given this inherent RFA limitation, we implemented an input randomization strategy (random reassignment of values to alleles) to minimize potential bias. For this, M random permutations of each gene’s allelic numbering in the original dataset is performed, effectively creating M independent input matrices. The RFA is run over the input matrices and in the main results we present each gene’s average MDA score. A sensitivity analysis was performed by comparing RFA results between two independent sets of *M* = 50 input matrices (effectively comparing 100 independent runs) (Fig. S5). Results suggest that the existing biases in independent runs of the RFA due to the assumption of numerical predictors are virtually mitigated with our input randomization strategy approach, specially for experiments classifying serotype.

## Acknowledgements

The authors acknowledge the sequence data kindly given by Angela Brueggemann and Andries van Tonder, and Richard Moxon for the valuable comments on a previous version of this manuscript.

## Additional Information

The authors declare no competing interests.

## Author contributions statement

This research was funded by the European Research Council under the European Union’s Seventh Framework Programme (FP7/2007-2013) / ERC grant agreement no. 268904 - DIVERSITY (www.erc.europa.eu; www.royalsociety.org). The funders had no role in study design, data collection and analysis, decision to publish, or preparation of the manuscript. JL and SG designed the study. JL performed the experiments and wrote the initial manuscript. JL, EWR, SG and MCJM revised the manuscript. JL, UO, SJP and CM curated the data and revised gene functionality.

## Supplementary Material

Supplementary Figures S1-7 described in a separate PDF file. The dataset herein used, in the format of an allelic matrix is made available in Table S1 in a separate spreadsheet. Functional description and literature review of top-scoring genes mentioned in the main text are in supplementary text within a separate PDF file.

